# Integrating spatial transcriptomics count data with Crescendo improves visualization and detection of spatial gene patterns

**DOI:** 10.1101/2024.03.07.583997

**Authors:** Nghia Millard, Jonathan H. Chen, Mukta G. Palshikar, Karin Pelka, Maxwell Spurrell, Colles Price, Jiang He, Nir Hacohen, Soumya Raychaudhuri, Ilya Korsunsky

## Abstract

Spatial transcriptomics allows for the analysis of a cell’s gene expression in the context of its physical location. With spatial transcriptomics data, investigators often want to find genes of interest whose spatial patterns are biologically relevant in multiple samples. However, due to confounding factors in spatial data that produce noise across samples, datasets, and technologies, it is challenging to visualize genes and their spatial patterns across samples. We present Crescendo, an integration algorithm that performs correction directly on gene expression counts to reduce variation from technical confounders. We first apply Crescendo to a 3-sample spatial transcriptomics mouse brain dataset to show how Crescendo enables accurate visualization of gene expression across these spatial transcriptomic samples. We then demonstrate Crescendo’s scalability by integrating a 16-sample immuno-oncology dataset of 7 million cells. Finally, we show that Crescendo can perform cross-technology integration by merging a colorectal cancer (CRC) scRNA-seq dataset with two CRC spatial transcriptomics samples. By transferring information between technologies, Crescendo can impute poorly expressed genes to improve detection of gene-gene colocalization, such as ligand-receptor interactions.

## Background

High dimensional single-cell technologies^1–3^ enable the discovery and characterization of cellular heterogeneity and potential function of important cell states^4–8^. Data from single-cell RNA sequencing (scRNA-seq) and recently emerging spatial transcriptomics platforms that enable spatially resolved single-cell transcriptional profiling^9–13^ can be used to examine expression patterns of individual genes. Identifying key genes is an essential part of defining cellular functions, building regulatory networks, and understanding cell-cell interactions^14–18^. In non-spatial scRNA-seq, we often visualize clusters of cells in a uniform manifold approximation and projection (UMAP)^19^, upon which we can overlay gene expression to identify cell-type-specific patterns. However, in spatial transcriptomics, we can go a step further and observe the expression of key genes in individual cells in the context of their real physical location. Furthermore, spatial gene expression can be used to identify potential cell-cell communication via consistent colocalization of two genes with ligand-receptor analysis^16–18,20–23^. However, understanding true spatial gene expression patterns is difficult because the measurements of many genes are sparse or not captured well^24,25^, confounded by technical factors^26–28^, or expressed in a cell type that does not group together in physical space which makes effective visualization challenging^29–31^.

We and others previously showed that for datasets affected by confounding factors such as differences in batches, samples, and datasets, it is extremely important to integrate cells across those batches or samples. However, most single-cell integration algorithms such as Harmony^32^, Seurat anchor integration^33^, and mutual nearest-neighbors (MNN)^34^ modify a lower-dimensional representation of gene expression, rather than directly correcting the genes themselves. To facilitate the visualization of gene expression and identification of spatial gene patterns spatial tissue slices, it is crucial to remove the effect of confounding factors and provide a way to impute sparse or poorly captured gene expression across slices for individual genes. To our knowledge, only the bulk RNA-seq algorithm ComBat-Seq^35^ is explicitly designed to correct individual gene counts, and no existing method performs both correction and imputation.

Here, we present Crescendo, a novel solution to single-cell integration of gene counts. Crescendo is designed to simultaneously correct systematic variation across datasets and impute low-expressed gene counts that result from technical confounders. Crescendo was designed to work directly on count data. In this manuscript, we focused on gene correction in the context of spatial transcriptomics, where it is critical to observe the expression of key genes in spatially defined individual cells, rather than in clusters of cells. First, we showed that Crescendo correction facilitated the tracking of 3-dimensional gene expression in spatial transcriptomics data containing three serial sections of a mouse brain^36^. To showcase Crescendo’s scalability, we then performed temporal computational benchmarks on a 16-sample, 7-million-cell immuno-oncology spatial transcriptomics dataset^37^. Then, in a more challenging scenario, we used Crescendo to integrate a scRNA-seq colorectal cancer (CRC) dataset^38^ with CRC spatial transcriptomics samples. Finally, in proof-of-principle analyses, we illustrated that Crescendo-corrected gene expression enabled the detection of spatial ligand-receptor interactions that were obscured by technical effects.

## Results

### Crescendo corrects technical variation in gene expression across spatial datasets

Here, we showcase Crescendo in the context of spatial transcriptomics data. Crescendo is an extension of the Harmony algorithm, which removes technical effects in a lower-dimensional representation of data (**Figure 1A**), such as principal components from principal components analysis (PCA). After Harmony fits linear models to PCA embeddings, Crescendo fits generalized linear models to gene expression counts (**Figure 1B**, **Methods**). Both Harmony and Crescendo assume that technical effects are cell-type-specific. The result of Crescendo is corrected gene counts that can facilitate visualization of a gene across tissue slices (denoted here as batch); in some cases, this may improve the ability to visualize and detect gene spatial patterns in a slice (**Figure 1C**). Importantly, Crescendo preserves counts in the output expression matrix, making the final output amenable to count-based downstream analyses, such as visualization, differential expression, and spatial pattern analyses.

**Figure 1:**
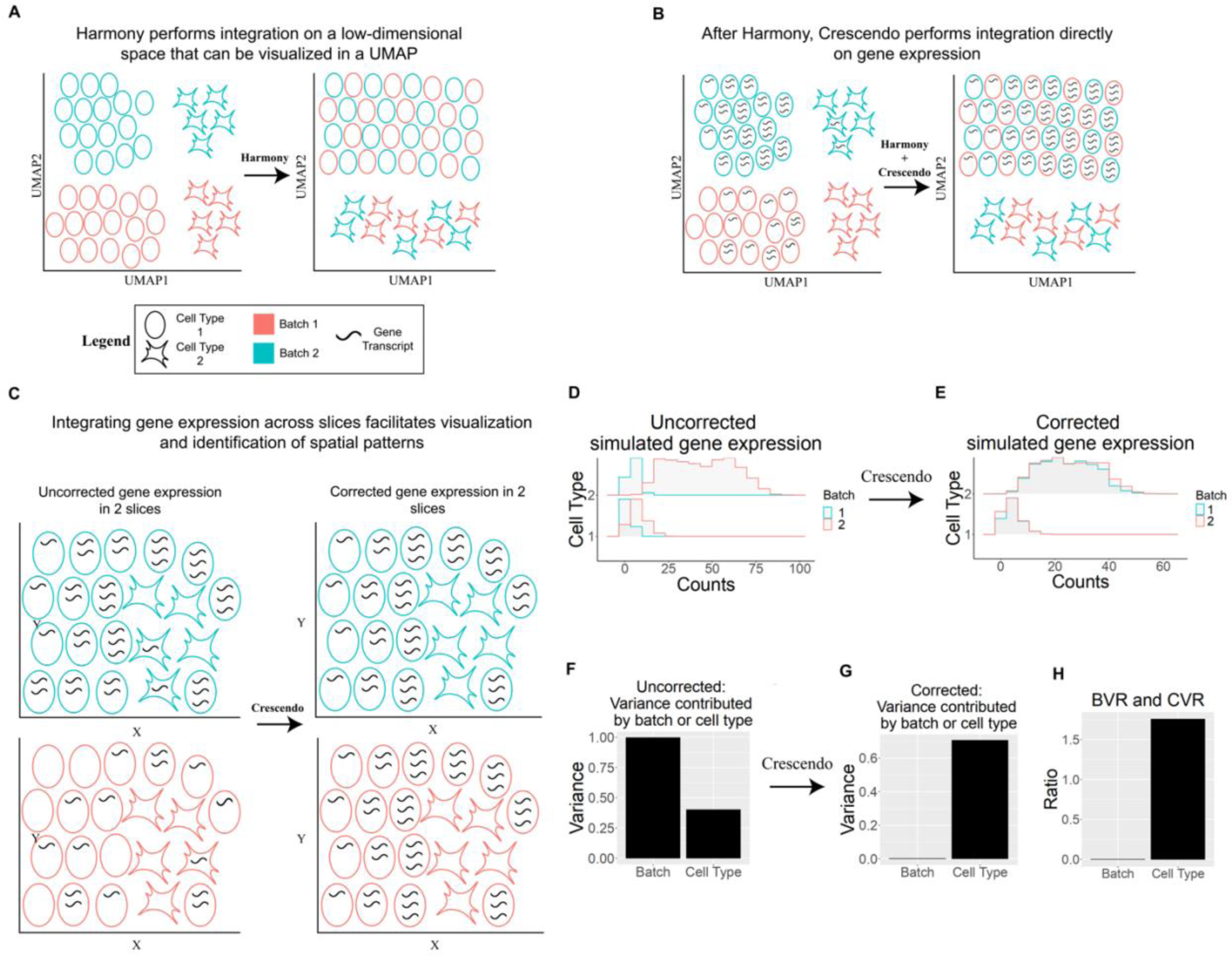
Crescendo directly corrects gene expression. **A,** Harmony integrates on lower-dimensional embeddings like principal components that are visualized with a 2-dimensional UMAP. **B,** Crescendo extends Harmony to correct gene expression, which can similarly be visualized in a UMAP. **C,** Spatial transcriptomics allows for visualization of gene expression in the context of cellular locations. Due to technical effects, gene expression can be poorly expressed and spatial patterns can be obscured. Crescendo infers the gene expression of a cell, which facilitates the visualization and spatial patten recognition of gene expression. **D,E**, Representative distributions of simulated gene expression before (**D**) and after (**E**) Crescendo correction. **F,G,** Batch-associated and cell-type-associated variance metrics before (**F**) correction and after (**G**) correction. **H,** Calculated batch-variance ratio and cell-type-variance ratio metrics based on **F-G**.

The inputs for Crescendo are a gene by counts matrix, cell-type information, and batch (or slice) information; the output is a corrected gene by counts matrix. To facilitate scalability, we allow users to first perform a biased downsampling to reduce the number of cells while accounting for rare cell states and batches; this is used for model fitting, but we still perform correction on all cells (**Supplementary Figure 1A, Methods**). After downsampling, we perform an estimation step in which we model how much variation in a gene’s expression derives from intrinsic biological sources (such as cell-type identity) and confounding technical sources (such as batch, sample, or technology). We then perform a marginalization step, in which we use the model from the estimation step to infer a model of gene expression without explicitly modeled confounding factors. Finally, we perform a matching step by using the original estimated model and the marginalized model to sample corrected counts (**Methods**). For lowly-expressed genes or those assayed with lower sensitivity, Crescendo can model gene expression assuming higher total read counts to perform imputation (**Methods**).

### Benchmarking gene-level correction with batch and cell-type variation metrics

Effective correction of gene expression must meet two objectives: (1) remove differences between cells that are driven by technical factors such as batch, sample, or technology and (2) preserve the biologically meaningful differences in gene expression, especially among cell types. To evaluate Crescendo, we developed two metrics to quantify the performance of gene expression correction: the batch-variance ratio (BVR) and cell-type-variance ratio (CVR). The first metric quantifies the removal of confounding factors (batch) as the ratio of batch-related variance in gene expression before correction versus after correction. Similarly, the second metric quantifies the preservation of cell-type-related differences as the ratio of cell-type-related variance in gene expression before versus after correction.

We calculate BVR and CVR by fitting generalized linear models in which we fit a gene’s counts with random effects for batch and user-defined cell-type identity (**Methods**). For each gene, we fit this model on both the uncorrected and corrected data. To obtain the BVR, we calculate the ratio of the batch-related variances between these fitted models; similarly, we calculate the CVR from the cell-type-related variances from these models. Ideally, correction will decrease variance associated with batch, which lowers the post-correction batch variance to give a BVR < 1. Furthermore, we ideally want to maintain or increase cell-type variance after correction, which would give a CVR >= 1; empirical observations from real data suggest that a CVR >= 0.5 is generally good preservation of cell-type variability. We also note that if batch variance is initially low, signaling that a gene is not significantly affected by confounding factors, correction may not be necessary.

To demonstrate these metrics, we show example genes that exhibit high or low BVRs/CVRs after we performed gene expression correction on 3 samples from the Vizgen mouse brain receptor map dataset^36^ with both Crescendo (**Supplementary Figure 1B-C**) and Seurat anchor integration^33^ (**Supplementary Figure D-E**). We then applied Crescendo on simulated gene expression data. To simulate a gene count distribution, we first simulated cells from different batches and cell types. We then simulated batch-specific and cell-type-specific gene expression rates to parameterize a Poisson distribution from which we sampled gene counts for each cell (**Figure 1D-E, Methods**). For this representative gene, we performed Crescendo integration and calculated the BVR and CVR metrics (**Figure 1F-H**). Over 10,000 gene simulations, we observed that Crescendo dramatically decreased technical noise in 100% of the simulated genes (BVR < 1), with 98.64% of those genes also exhibiting CVR >= 0.5 (**Supplementary Figure 1F**).

### Crescendo corrects technical effects across serial sections in whole mouse brain

We then designed an analysis to demonstrate the practical utility of Crescendo to correct gene expression, and by doing so, improve visualization of gene expression in physical space. We used a public spatial transcriptomics dataset of the mouse brain profiled by the Vizgen MERSCOPE platform^36^. We performed a standard scRNA-seq analysis pipeline^39^ to analyze and cluster three serial coronal slices (S3R1, S3R2, S3R3) from the same mouse brain that represent batches; in aggregate, this data contains *in situ* expression for 483 genes in 179,385 segmented cells (**Figure 2A**). This dataset features inhibitory and excitatory neuronal subtypes, along with astrocytes, microglia, oligodendrocyte progenitor cells (OPCs), and endothelial cells (**Figure 2B**). Technical effects were variable, with certain cell types (*e.g.,* inhibitory and excitatory neuronal subtypes) exhibiting greater levels of technical effects than others (**Supplementary Figure 2A**). In physical space, neurons tended to be well-organized, while cell types such as astrocytes and microglia were dispersed across the sections (**Figure 2C, Supplementary Figure 2B**).

**Figure 2:**
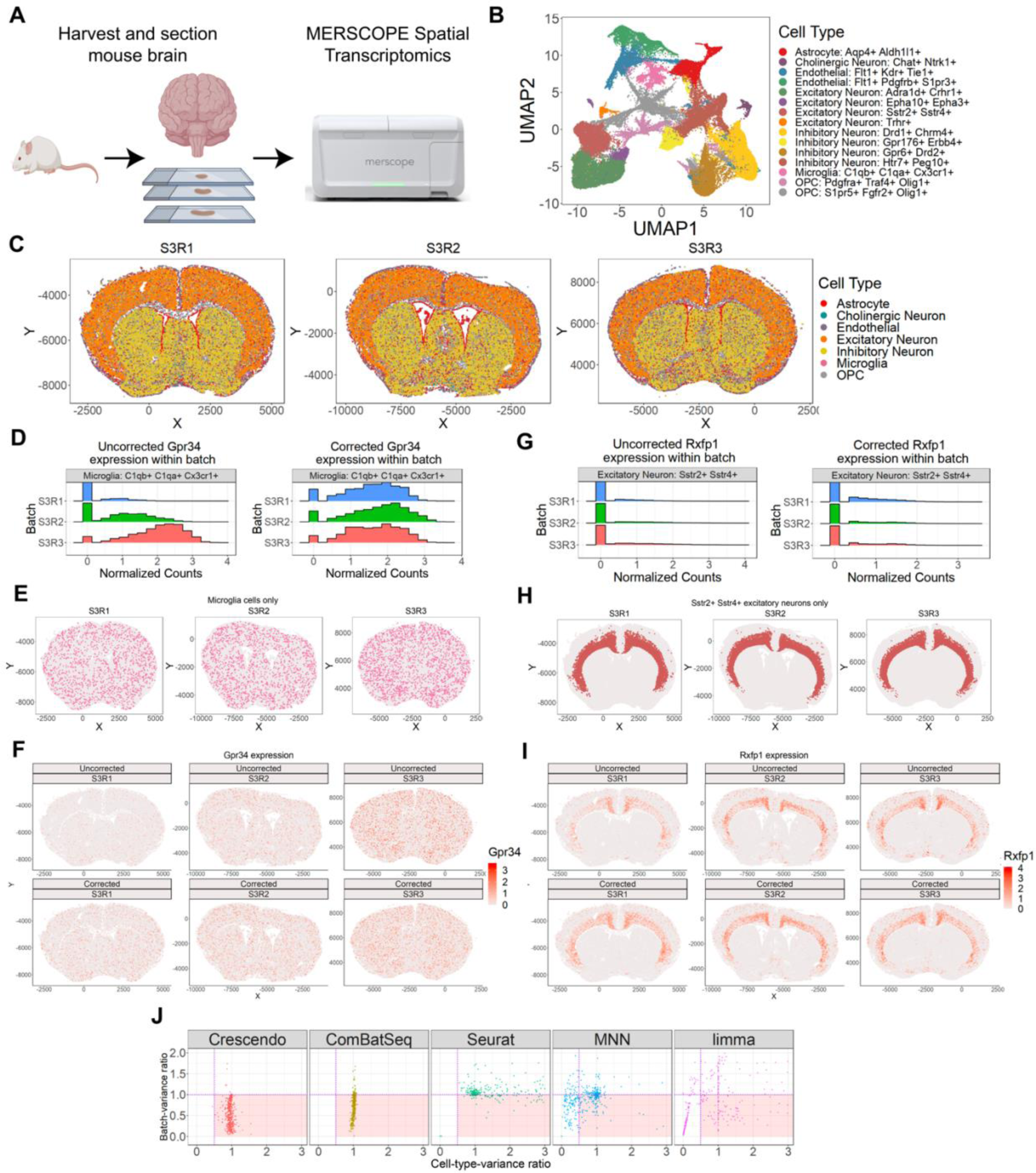
Crescendo facilitates visualization of genes across spatial transcriptomics datasets of serial sections from whole mouse brain tissue. **A,** The Vizgen MERSCOPE platform was used to assay three coronal mouse brain tissue slices^36^. **B,** Cell state classifications of cells based on marker genes. **C,** Spatial locations of broad cell types. **D,G,** Gene expression distributions across slices for *Gpr34* (**D**) and *Rxfp1* (**G**). **E,H,** Spatial locations of cell types with the highest expression *Gpr34* (**E**) and *Rxfp1* (**H**). **F, I** Gene expression visualizations in physical space before and after Crescendo correction for *Gpr34* (**F**) and *Rxfp1* (**I**). **J,** Scatter plots of batch-variance ratio (BVR) and cell-type-variance ratio (CVR) metrics calculated for all 483 genes across 5 different integration algorithms. Purple dashed vertical line is at CVR = 0.5 and the purple dashed horizontal line is at BVR = 1. Red at BVR < 1 and CVR >= 0.5 is the target zone for genes that were corrected well.

Because these slices represent a z-stack of serial sections in a similar area of the brain, we expected genes to be expressed at consistent levels across the slices (denoted as batch). However, we observed that several genes exhibited noticeable batch-related variance, though it tended to be smaller in magnitude compared to cell-type variance (**Supplementary Figure 3A**). To begin, we analyzed the effect of Crescendo on three genes that were cell-type specific: *Gpr34* in microglia cells^40,41^, *Rxfp1*^42,43^ in cortical excitatory neurons, and *Epha8*^44^ in striatal inhibitory neurons. Each of these genes were subject to technical effects (**Figure 2D, 2G**, **Supplementary Figure 4A-E**). For each gene, we show that correction improves visualization for a gene by making expression more consistent across the slices.

We first looked at the gene *Gpr34*, which is predominantly expressed by microglia, a cell type that clustered together tightly in the UMAP (**Figure 2B, Supplementary Figure 4A**). However, in physical space, both microglia (**Figure 2E**) and the expression of *Gpr34* (**Figure 2F**) are spread out across the slices, making visualization challenging even when plotting *Gpr34*-expressing cells on top. This visualization is even worse in slice S3R1, which has overall lower expression (**Figure 2D,2F**). After using Crescendo to correct *Gpr34* expression, we observed noticeably higher expression of *Gpr34* in S3R1 at levels relative to the other two slices, and more even expression across all slices (**Figure 2D, 2F)**.

We next looked at the gene *Rxfp1*, which is predominantly expressed by *Sstr2+ Sstr4+* excitatory neurons (**Figure 2B, Supplementary Figure 4B**). Here, we observed *Rxfp1* expression at similar maximal levels across all slices but noticed that many *Sstr2+ Sstr4+* excitatory neurons in S3R1 had noticeably lower levels of *Rxfp1* expression (**Figure 2G-I**). In physical space, we observed that *Sstr2+ Sstr4+* excitatory neurons tended to cluster in specific layers of the cortex (**Figure 2H**), but visualization of *Rxfp1* expression showed that expression was not consistent across these neurons in the same tissue (**Figure 2I**). Again, after using Crescendo to correct *Rxfp1* expression, we observed more even expression across all slices while importantly not increasing expression in other cell types such as excitatory neurons in the other cortical layers (**Figure 2G, 2I, Supplementary Figure 5A-B**).

Finally, we looked at the gene *Epha8*, which is predominantly expressed by some inhibitory neuron states (**Figure 2B, Supplementary Figure 4C**). *Epha8* expression was also subject to technical effects, with low expression in slice S3R3 (**Supplementary Figure 4C-E**). After correction with Crescendo, we were again able to observe relatively even *Epha8* expression across all slices (**Supplementary Figure 4D-E**).

Our analyses so far demonstrate the utility of Crescendo to improve gene visualization by ameliorating technical effects in three genes. We next quantified how well Crescendo removes technical effects while retaining biological variation in all 483 genes in the MERFISH panel. We applied Crescendo to each gene and calculated the BVR and CVR metrics (**Figure 2J**). Of the 483 genes, Crescendo produced a BVR < 1, CVR >= 0.5 in 408 genes. We next compared Crescendo’s ability to correct individual genes in this mouse brain spatial dataset to four representative state-of-the-art algorithms: ComBat-Seq, Seurat anchor integration, MNN, and limma (**Methods**). With the caveat that these methods were not explicitly intended to be used for this purpose, we observed that Seurat, MNN, and limma struggled to remove batch-associated variation from gene expression (sometimes even increasing batch variation) or resulted in a dramatic decrease in biological cell-type variation (**Figure 2J**). Their poor performance is likely due to their assumptions of Gaussian structure in data rather than the count-based structure of gene expression. Of the 483 genes, ComBat-Seq, Seurat, MNN, and limma produced a BVR < 1, CVR >= 0.5 in 364, 142, 160, and 104 genes, respectively, compared to Crescendo’s 408 genes.

### Crescendo scales efficiently to millions of cells

Single-cell datasets are increasing in size, with experiments regularly profiling 100,000+ cells per experiment, the creation of large single-cell atlases on the order of millions of cells, and spatial transcriptomics experiments potentially profiling 100,000+ cells per slice^45,46^. This leap in data size makes computational efficiency critical for gene-level correction. We tested the ability of Crescendo and other methods to scale to both many cells and many tissues by using an Immuno-oncology FFPE dataset produced by the Vizgen MERSCOPE platform^37^. The Immuno-oncology dataset features a custom 500-gene panel designed to profile immune, stromal, and malignant cells across 9 different tissue types. This collection contains 7,020,548 post-QC cells across 16 individual slices spanning 8 tissue types (**Methods**).

We first attempted to correct all 500 genes with each method (**Supplementary Figure 6A**). ComBat-Seq, Seurat, and MNN failed to complete due to memory requirements while limma took 6.6 hours. Unlike alternative methods, Crescendo fits and corrects a gene independent of others, which allows users to fit genes individually and reduces the risk of running into memory complications. Overall, Crescendo performed best by correcting all 500 genes in 3.3 hours. For the sake of comparison, we also summed processing time across genes to simulate downsampling each gene individually (though this is unnecessary, **Methods**), which took 6.1 hours.

In contrast with the alternative methods that use the information from all genes (removing a gene changes results), Crescendo allows users to correct genes independently and prioritize specific genes of interest. To better understand the scaling behavior of Crescendo based on the number of genes and cells corrected, we repeatedly corrected random samples of genes in increasingly larger subsamples of the 7 million cells. In each run, we corrected 1, 2, 5, 10, or 50 random genes 100 different times for each subsample of 10K, 25K, 100K, 250K, 500K, 750K, 1M, 2M, 3M, 4M, 5M, 6M, or all 7M cells (**Methods**). Impressively, Crescendo was able to consistently correct 50 genes across 7 million cells in less than 7 minutes (**Supplementary Figure 6B**. For each run, we also isolated the amount of time Crescendo takes to perform the downsampling, estimation, marginalization, and matching steps (**Supplementary Figure 6C**). Computational runtime for the downsampling step was dependent on the number of cells while runtime for the estimation step scaled relatively linearly based on the number of genes being corrected. The marginalization and matching steps also depended on the number of genes but scaled slightly better than linearly.

We also performed singular runs of correcting all 500 genes at once for each dataset size. For all 500 genes at once, Crescendo was able to process a 3 million-cell dataset in ∼22 minutes (**Supplementary Figure 6D**); the estimation step ran into memory issues when fitting GLMs on dataset sizes of greater than 4 million.

### Crescendo corrects technology effects by integrating paired colorectal cancer scRNA-seq and spatial transcriptomics datasets

We hypothesized that Crescendo could be used to integrate and impute gene expression between non-spatial technologies, which captures the full transcriptome, and spatial technologies, which gives physical locations of transcripts. To demonstrate this, we used Crescendo to correct technology effects by integrating two human colorectal cancer (CRC) spatial transcriptomics slices with a CRC 10X scRNA-seq dataset^38^. The scRNA-seq dataset contains 69,153 cells from 29 CRC tissue samples while the two spatial transcriptomics slices (PFA_A6 and PFA_A11) were both generated from the same CRC tissue sample taken from a donor in the scRNA-seq dataset (**Figure 3A, Methods**). The two technologies share 477 common genes, reflecting the small gene panels currently available to spatial transcriptomics datasets. Integrating these two technologies represents a more challenging scenario because the technology effects are noticeably larger than the batch effects between the spatial transcriptomics slices (**Figure 3B, Methods**). Human CRC tissue also contains much less organization than the highly structured quality of the brain, further complicating gene visualization. Moreover, cells in primary human tissue tend to be packed close to each other, making overplotting even more problematic. To address this, we plotted gene expression similar to the mouse brain section by plotting gene-expressing cells over non-expressing cells to represent a best-case scenario of visualizing gene expression across slices.

**Figure 3:**
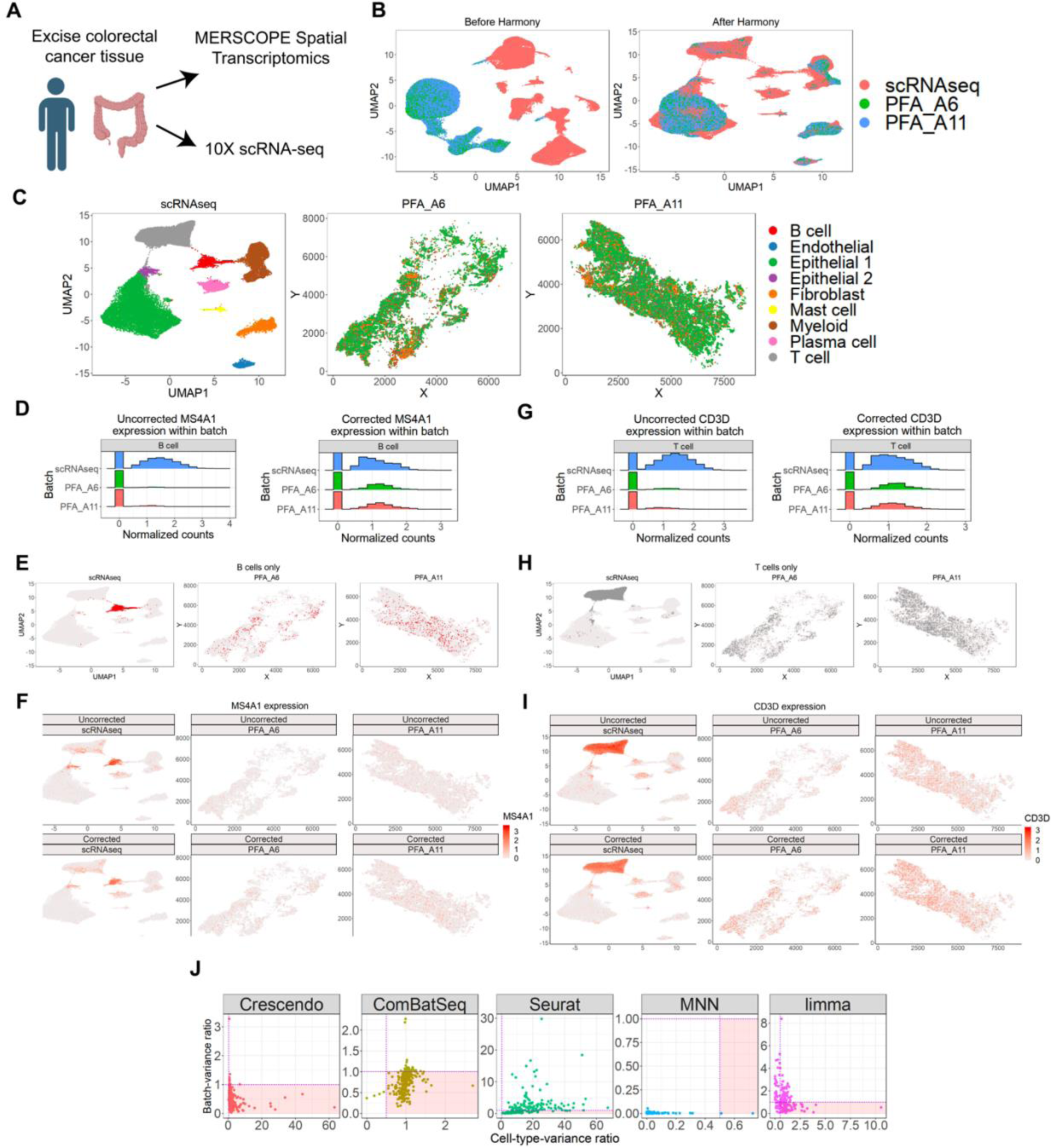
Crescendo corrects technology effects between a colorectal cancer (CRC) scRNA-seq dataset and two CRC spatial transcriptomics samples. **A,** Colorectal cancer samples were assayed with scRNA-seq and spatial transcriptomics. These datasets shared 477 genes **B,** UMAP embedding of cells from scRNA-seq and spatial transcriptomics before and after batch integration with Harmony (integration performed on a dataset variable where the scRNA-seq dataset and each spatial slice was considered a dataset). **C,** Broad cell type classification of cells and spatial locations of cell types in spatial slices (middle, right). **D,G,** Gene expression distributions across slices for *MS4A1* (**D**) and *CD3D* (**G**). **E,H**, In these and following plots, scRNA-seq is plotted in UMAP space, while spatial slices are plotted in physical space. Spatial locations of cell types with the highest expression of *MS4A1* (**E**) and *CD3D* (**H**). **F, I** Gene expression visualizations in physical space before and after Crescendo correction for *MS4A1* (**F**) and *CD3D* (**I**). **J,** Scatter plots of batch-variance ratio (BVR) and cell-type-variance ratio (CVR) metrics calculated for all 477 genes across 5 different integration algorithms. Purple dashed vertical line is at CVR = 0.5 and the purple dashed horizontal line is at BVR = 1. Red at BVR < 1 and CVR >= 0.5 is the target zone for genes that were corrected well.

Similar to the author-defined cell types in the scRNA-seq data, we identified epithelial cancer cells, fibroblasts, endothelial cells, T cells, B cells, plasma cells, and myeloid cells in the spatial transcriptomics slices (**Figure 3C**; **Methods**). In physical space, all cell types were relatively spread out, though some cell types such as epithelial cells and fibroblasts occasionally formed small aggregates.

After integrating the scRNA-seq data and spatial slices together with Harmony (**Figure 3B, Methods**), we compared gene expression between the spatial slices and the scRNA-seq data; we observed several genes that were expressed in most cells of a cell type in the scRNA-seq data but not well-expressed in that same cell type in the spatial transcriptomics slices. For instance, the gene *MS4A1* (*CD20*) is a marker for B cells and was well-expressed in the scRNA-seq data but was not expressed well in the spatial slices (**Figure 3D**). After Crescendo correction, we observed increased expression of *MS4A1* in the spatial slices on a level more similar to the scRNA-seq data; this also provided easier visualization of *MS4A1* expression in the spatial slices that was consistent with the locations of B cells (**Figure 3D-F**). We observed similar trends for the T-cell-specific gene *CD3D*. Unlike *MS4A1*, expression of *CD3D* was visible in physical space, but the level of expression is still lower than scRNA-seq (**Figure 3I**). After correction, we observed more even *CD3D* expression across all three datasets and strengthened *CD3D* expression, particularly in PFA_A6, in the spatial slices that was consistent with the locations of T cells (**Figure 3G-I**).

Subsequently, we performed correction and calculated the BVR and CVR metrics on all 477 genes in the CRC scRNA-seq and spatial datasets with Crescendo and the other 4 benchmarking methods (**Methods**). We observed that of the 477 genes, 439 exhibited a batch variance greater than 0.001 (**Supplementary Figure 3B, Methods**). Of these 439 genes, Crescendo, ComBat-Seq, Seurat, MNN, and limma provided a BVR < 1, CVR >= 0.5 in 423, 372, 78, 2, and 89 genes, respectively (**Figure 3J**). We had to plot on different scales since the ranges of BVR and CVR varied widely by method, with Seurat having the notably extreme maximum BVR of 30.

### Integrating spatial transcriptomics gene expression facilitates the identification of spatial ligand-receptor interactions via gene-gene correlations

We next looked at the spatial patterns of gene expression in physical space. A powerful aspect of single-cell spatial transcriptomics is the ability to simultaneously look at gene expression of a cell in the context of its physical neighbors. This view lets us hypothesize about potential interactions between neighboring cells through gene-gene interactions, particularly ligand-receptor interactions. To evaluate the ability of Crescendo to inform and improve the power to detect gene-gene interactions, we analyzed the correlation between a cell’s gene expression with that of its spatially neighboring cells.

Many investigators want to find spatial patterns between genes in specific cell types of interest. Thus, in these analyses, we calculated a spatial cross-correlation index (SCI) between genes in a cell-type-aware manner (**Methods**). Briefly, we subsetted cells within a slice to two cell types; for cell-type 1, we identified its nearest neighbors within a 30μm Euclidean distance that were from cell-type 2 and calculated their average gene expression to obtain a nearest-neighbor expression matrix. We then correlated the cell-type 1 gene expression matrix with the nearest-neighbor expression matrix (that encodes their cell-type 2 neighbors). We repeated this procedure for all combinations of cell types in each slice independently. Two genes in two different cell types that share similar spatial expression patterns exhibit a positive SCI, dissimilar patterns exhibit a negative SCI, and an SCI of zero indicates no consistent spatial pattern between the genes (**Figure 4A**).

**Figure 4:**
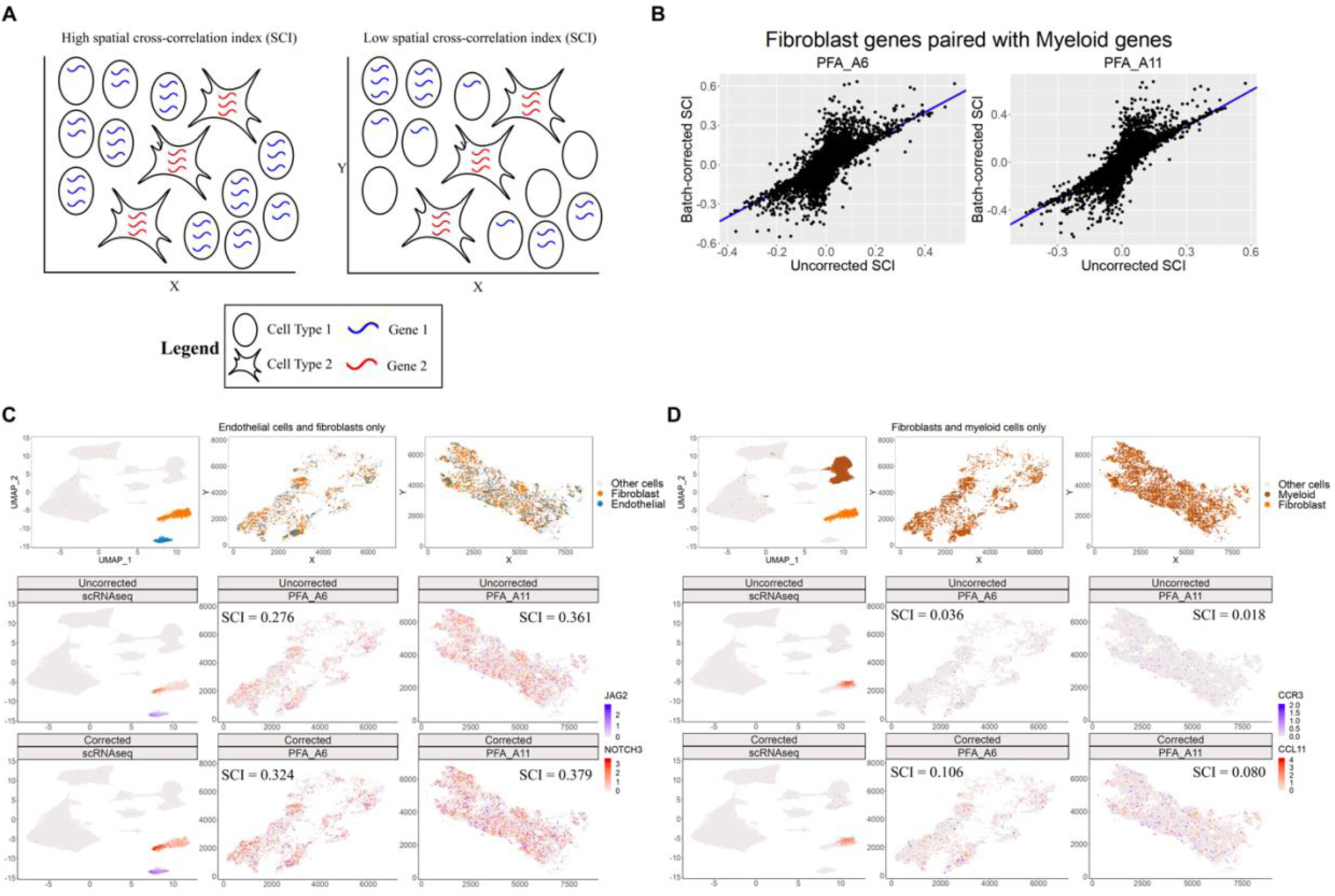
Crescendo correction increases ability to visualize and detect spatial gene-gene correlations. **A,** Example schematics of gene-gene pairs that have a high spatial cross-correlation index (SCI) and a low SCI. **B,** Comparison of SCIs for all fibroblast and myeloid cell gene-gene pairs before correction vs. after correction. **C,** In these and following plots, scRNA-seq is plotted in UMAP space, while spatial slices are plotted in physical space. Spatial locations of fibroblasts and endothelial cells (top). Gene expression visualization of *JAG2* in endothelial cells and *NOTCH3* in fibroblasts (bottom). SCIs are listed for each spatial sample before and after correction. **D,** Spatial locations of myeloid cells and fibroblasts (top). Gene expression visualization of *CCR3* in myeloid cells and *CCL11* in fibroblasts (bottom). SCIs are listed for each spatial sample before and after correction.

Overall, we observed that for a majority of gene-gene pairs in a cell-type pair, the SCI did not noticeably change after correction; however, we did observe several pairs that had a low uncorrected SCI change to a higher corrected SCI (**Figure 4B**). We chose two such examples that are well-studied ligand-receptor pairs to observe how Crescendo correction affects both visualization and SCI.

First, we looked at how *JAG2* expression in endothelial cells and *NOTCH3* expression in fibroblasts formed spatial coherent patterns. Previous studies^47^ show that *NOTCH3* signaling can drive transcriptional and spatial gradients in fibroblasts after interacting with Notch ligands, like Jagged-2, from vascular endothelial cells. In physical space, we observed areas of colocalization between endothelial cells and fibroblasts, and that the SCI for *JAG2* in endothelial cells and *NOTCH3* in fibroblasts was initially 0.276 in PFA_A6 and 0.361 in PFA_A11 (**Figure 4C**). However, *JAG2* expression in some cells was difficult to visualize due to technical effects. After correction with Crescendo, we observed more visible expression of *JAG2* in endothelial cells in both slices, which made identification of colocalizing *JAG2*-expressing endothelial cells and *NOTCH3*-expressing fibroblasts easier (**Figure 4C**). Statistically, the SCI increased to 0.324 in PFA_A6 and 0.379 in PFA_A11.

Next, we looked at *CCR3* expression in myeloid cells and expression of its ligand *CCL11* in fibroblasts, involved in the chemotaxis of leukocytes^48,49^ (**Figure 4D**). Notably, we observed that *CCR3* was mainly expressed by a subset of myeloid cells, with much better expression in the spatial transcriptomics slices. Conversely, *CCL11* expression was much higher in the scRNA-seq dataset. With technology-specific low expression of both genes, it was perhaps unsurprising that we saw low SCIs of 0.036 in PFA_A6 and 0.018 in PFA_A11, suggesting almost non-existent colocalization of these genes (**Figure 4D**). However, after correction with Crescendo, visualization of this gene-gene pair showed a modest increase in *CCR3* expression in some myeloid cells and a dramatic increase in *CCL11* expression such that areas where these genes colocalize are now visible (**Figure 4D**). Statistically, SCI noticeably increased to 0.106 in PFA_A6 and to 0.080 in PFA_A11. We note that the lower SCI values for this gene-gene pair is due to colocalization of these genes’ expression being limited to certain areas of the tissue while the *JAG2-NOTCH3* pair was more ubiquitously expressed within the specified cell types. Overall, these results suggest that Crescendo can help recover spatial patterns that were previously obscured by technical effects.

We then looked at *COL1A2* expression in fibroblasts and *CXCL14* in myeloid cells, which have no previously known interactions. *CXCL14* expression was noticeably low in both the spatial datasets and the scRNA-seq while *COL1A2* was well-expressed primarily in the scRNA-seq dataset. With such low expression in both genes in the spatial datasets, the SCI was notably low in both slices: −0.009 for PFA_A6 and −0.004 for PFA_A11 (**Supplementary Figure 7A-B**). After correction, we observed that *COL1A2* expression was noticeably increased in the spatial datasets but *CXCL14* was still low; this resulted in a dramatically decreased SCI in both slices to −0.455 in PFA_A6 and −0.506 in PFA_A11 (**Supplementary Figure 7B**).

Finally, we reasoned that if two cell types colocalize, then the SCI between their markers should be relatively high. Fibroblasts and T cells are abundant cell types in the spatial slices and visually appear close to each other in many areas (**Supplementary Figure 7C**). However, the SCI between a pair of their markers, *FN1* in fibroblasts and *CD3E* in T cells, was relatively low at 0.024 in the first slice and 0.016 in the second (**Supplementary Figure 7D)**. Visualization of these marker genes showed that the low spatial cross-correlation is explained by the low expression of these genes. After correction with Crescendo, we observed much more visible expression of both *FN1* and *CD3E* in both slices (**Supplementary Figure 7D)** with an accompanying increase in SCI to 0.248 in the first slice and to 0.290 in the second slice.

## Discussion

Identifying genes or features of interest is an important aspect of generating hypotheses from single-cell data. In spatial transcriptomics data, visualizing a gene’s spatial patterns can help infer the role of a gene in the function of a cell type and the localization of cell types to specific niches. Thus, it is important to correct and impute gene expression in order to accurately visualize it. Here, we introduced Crescendo, which accepts Harmony outputs and gene counts as input and returns corrected counts as output. We showed that Crescendo can remove technical effects from a vast majority of genes in spatial transcriptomics data, which facilitated better visualization of the gene’s expression across tissue slices and overall gene spatial patterns. We also developed statistics, BVR and CVR, to quantify the performance of correcting genes and demonstrated that Crescendo outperforms alternative methods.

Crescendo is scalable to millions of cells, which enables it to accommodate the large number of cells featured in modern single-cell spatial transcriptomic datasets^50–53^ and single-cell atlases^7,54,55^. We showcased Crescendo’s scalability by correcting genes in 7 million cells across 16 batches. We predict that spatial datasets will continually grow to incorporate more individuals and multiple samples from the same individual, thus making scalability even more important.

We observed that identifying gene-gene interactions can be difficult if expression of one or both genes is too low. In many examples we showed, gene expression was good in at least one dataset, which helped rescue expression in the other datasets. However, if a gene is lowly expressed across all datasets, then correction and/or imputation will not rescue its expression. Poor expression of certain genes like cytokines can be observed in many technologies^24,25,56–59^, so investigators should consider how well a gene is expressed before attempting to use Crescendo correction to impute gene expression.

Accurately correcting technical effects in fluorescence in situ hybridization (FISH)-based spatial datasets is challenging because cell segmentation is a significant challenge^31,60–62^. Inaccurate segmentation can erroneously assign certain transcripts to the wrong cell. Since the number of unique genes expressed and the transcripts per cell in spatial data tends to be significantly lower than scRNA-seq^25,61^, erroneous assignment of transcripts to a cell makes the data considerably noisier and accurate correction of genes more difficult. Indeed, in all spatial datasets we showcased, we observed several instances of a cell type containing transcripts of markers for other cell types (*e.g.,* a B cell marker in T cells). Theoretically, segmentation could cause systematic errors in transcript assignments; for example, if B cells and T cells tend to colocalize, there is a higher chance for their transcripts to be erroneously assigned amongst each other. If transcripts are systematically erroneously assigned, it is possible that correction and imputation may increase expression of an erroneous gene. We speculate that as segmentation performance increases, the effectiveness and accuracy of Crescendo correction should increase as well.

The Crescendo framework has other potential applications because it models counts, which are present in data generated from other technologies. For example, Crescendo could theoretically correct counts for single-cell ATAC-seq data^63–65^ and genomic data. Integrating genomic counts may also be useful for quantitative trait loci (QTL) analyses^66–68^ if they are confounded by technical noise. Due to the visual benefits of correcting gene counts to be more even across datasets (or tissue slices), we envision that investigators will utilize Crescendo to aid in gene visualization and hypothesis generation in scRNA-seq or spatial transcriptomics datasets.

## Supplementary Figure Legends

**Supplementary Figure 1: Downsampling and simulation results from Crescendo. A,** Comparison between model estimates for a gene when fit on full or downsampled data – outlier is the intercept, while other values are variable coefficients. **B,** Example gene distribution for a gene that exhibits a low (good) BVR after correction. **C,** Example gene distribution for a gene that exhibits a high CVR (good) after correction, where only the proper cell type (microglia) is corrected, and no other cell types gain expression. **D,** Example gene distribution for a gene that exhibits a high (bad) BVR after correction, where batch effect actually increases. **E,** Example gene distribution for a gene that exhibits a low CVR (bad) after correction, where other cell types erroneously gain expression of a gene and reduces the marker potential of a gene. **EF,** X-Y plots of BVR and CVR metrics calculated for all 10,000 simulated genes when correcting with Crescendo. Purple dashed vertical line is at 0.5 and the purple dashed horizontal line is at 1. Red shaded box encompasses all points with BVR < 1 and CVR >= 0.5, the target zone for genes that were corrected well.

**Supplementary Figure 2: Technical effects in the mouse brain coronal sections and cell type spatial locations. A,** UMAP of cells from all three sections before and after Harmony integration. **B**, Spatial locations of cell types in the S3R1 slice; because the other slices are serial sections, the locations of most cell types are similar in the other slices.

**Supplementary Figure 3: Cell-type variance and batch variance contributions for each gene. A,** Each dot depicts a gene. This plot compares the batch variance and cell-type variance for each gene from the mouse brain dataset. Purple dashed line has a slope of 1. **B,** Each dot depicts a gene. This plot compares the batch variance and cell-type variance for each gene from the integrated colorectal cancer (CRC) scRNA-seq dataset and the two CRC spatial transcriptomic slices. Purple dashed line has a slope of 1.

**Supplementary Figure 4: Mouse brain dataset gene expression in UMAP space and relevant visualizations of the gene *Epha8 and Rxfp1*. A-C,** Gene expression visualizations of *Gpr34*, *Rxfp1*, or *Epha8* in UMAP space before and after gene expression correction with Crescendo. **D,** Gene expression distributions of *Epha8* across slices. **E,** Spatial locations of *Epha8*-expressing cells and gene expression visualizations in physical space before and after Crescendo correction.

**Supplementary Figure 5: Crescendo scales to millions of cells. A,** Total runtime for correcting 500 genes in 7 million cells across 5 integration algorithms. Crescendo was run on genes individually, either downsampling once (Crescendo_Down1) or for downsampling independently for each gene (Crescendo_DownAll). Methods labeled “Failed to finish” failed due to computational memory restraints. **B**, Total runtime for Crescendo across a range of dataset sizes and number of genes. Genes were randomly selected for correction; each pair of dataset size and gene number was run 100 times to estimate the average time it takes to correct. Points denote the average runtime and error bars denote standard deviation. **C,** Runtimes for the downsampling, estimation, marginalization, and matching steps for the runs described in panel **B**. **D,** Runtimes for running Crescendo on all 500 genes at once.

**Supplementary Figure 6: Visualizations of the gene *TRAC* in the integrated CRC scRNA-seq and spatial datasets. A,** Gene expression distributions of *TRAC* across the scRNA-seq dataset and the two spatial transcriptomic slices. **B,** Spatial locations of T cells. **C,** gene expression visualization of *TRAC* in physical space before and after Crescendo correction.

**Supplementary Figure 7: Crescendo correction improves gene expression visualization and uncovers gene spatial patterns. A,** In these and following plots, scRNA-seq is plotted in UMAP space, while spatial slices are plotted in physical space. Spatial locations of myeloid cells and fibroblasts. **B,** Gene expression visualization of *COL1A2* in fibroblasts and *CXCL14* in myeloid cells. SCIs are listed for each spatial sample before and after correction. **C,** Spatial locations of fibroblasts and T cells. **D,** Gene expression visualization of *FN1* in fibroblasts and *CD3E* in T cells. SCIs are listed for each spatial sample before and after correction.

## Methods

### 1. Crescendo

#### 1.1 Overview

The Crescendo algorithm extends the Harmony algorithm to perform integration on gene count distributions. Harmony performs integration on a lower-dimensional latent space such as principal components and assumes the input data is Gaussian. In brief, Harmony performs iterative soft clustering to group similar cells across batches/samples/datasets and calculates cell-type-specific correction factors. Crescendo is applied to genes individually. It takes as input the count of that gene (x_i_) and their batch assignments (ϕ_i_) for each cell *i*. Crescendo also utilizes Harmony cluster soft-cluster assignments (*R_i,j_*) which is a number between 0 and 1 representing the probabilistic membership of cell *i* in cluster *j* (though any cluster assignment is theoretically compatible with the Crescendo framework). The output of Crescendo is a corrected gene count (x_i_*) for each cell *i*. After integrating cells from multiple batches into a shared low dimensional embedding with the Harmony algorithm, Crescendo builds an ‘observed’ Poisson generalized linear mixture model (GLMM) of a gene that includes both latent biological (cell-type) and batch variation. To infer a batch-free model, Crescendo analytically integrates this GLMM. The observed and batch-free models are each used to parameterize Poisson distributions that represent probability distributions for observing a specific count. Finally, we utilize inverse cumulative distribution function (CDF) mapping to sample a new batch-free count for each cell under the batch-free model that has an equal likelihood to the observed count under the observed model. Implementations of Crescendo are available as part of an R package at https://github.cmo/immunogenomics/crescendo.

We chose to use the Poisson distribution instead of a Gaussian linear model, as linear models tend to produce inflated corrected counts; this is because many observed gene counts are 0. Count-based distributions such as Poisson and Negative Binomial more accurately fit the count-based nature of single-cell data. For most genes, we observe that gene expression within a cell-type and within a batch tends to be poisson distributed, even if the overall gene expression distribution is more complex. As a result, we fit Poisson GLMMs.

#### 1.2 Fitting Poisson gene count models

To build an intuition for Crescendo, we first start with modeling and removing batch effects for one gene in one cell type.

To fit a Poisson generalized linear model for a gene, we fit the counts for cell *i* as the dependent variable in order to obtain a rate (**μ**_i_). By default, we fit latent biological (cell-type) and batch terms, in addition to fitting the total number of unique molecular identifiers per cell (nUMI) as an offset. We interpret latent biological variation as wanted variation (X_i_), while we interpret batch variation as unwanted variation (Y_i_). For simplicity, we assume that X and Y are independent. We explicitly fit unwanted variation variables (Y_i_) as random effects, which model coefficients as a normal distribution centered at 0 with an empirically derived variance. Potential candidate variables for unwanted variation include different batches, samples, datasets, technologies, and fields-of-view (spatial). We note that investigators may want to keep certain the variation that derives from intended experimental categories, such as the variation from stimulations or time points; such variables would be included as wanted variation (X_i_) instead. Overall for a gene, we fit the following <u>observed</u> model:

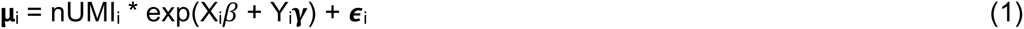

Where *β* and **γ** represent fitted coefficients for the wanted and unwanted variation terms respectively, and ***ϵ*** represents additive residuals. Since unwanted variation terms are fitted as a random effect, it follows that for *M* unwanted variation terms their distribution is:

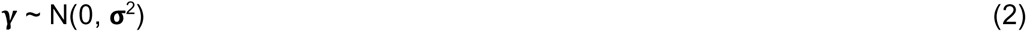

Where sigma is an empirically-derived variance from fitting the GLMM.

Equation (1) is equivalent to:

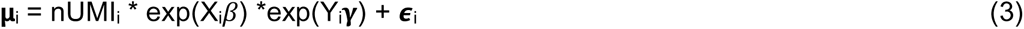

Fitting equation (3) gives the <u>observed</u> model. For each cell, the left-hand-side (LHS) of equation (3) is based on their batch and cell-type identity, which is then used in the marginalization and matching steps. To fit equation (3), we used the “glmnet” function from the R package “glmnet” with parameters alpha = 0, lambda = NULL, family = “poisson”. We take the first column of coefficients from the results as the estimated gene expression coefficients (right-hand-side of equation 3).

#### 1.3 Correcting for batch effects with marginalization

We frame “correcting” for a categorical batch variable as marginalizing over that variable. Since we want to marginalize the effect of unwanted variation (Y_i_), we take the expectation over Y. Taking the expectation of equation 3 gives:

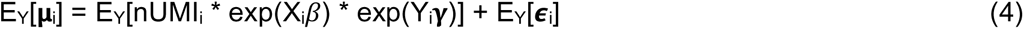

Since nUMI is a constant and we assumed that X and Y are independent, equation (4) simplifies to:

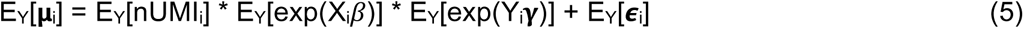

Only the third term depends on Y, so the expectations of the other terms are the terms themselves:

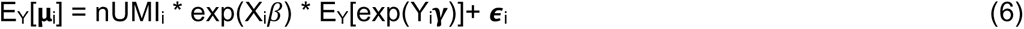

From equation 2, we see that exp(**γ**) yields a lognormal distribution with a known expectation, which allows us to analytically calculate the expectation:

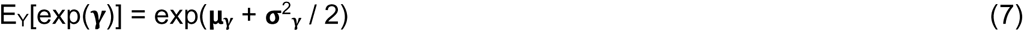

Furthermore, since we fit modeled **γ** as a random effect, **μ**_**γ**_= 0. With this and substituting equation (7) into equation (6), we obtain a final marginalized model:

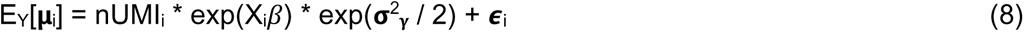

Which simplifies into the following linear transformation:

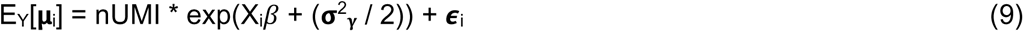

Let **μ**_i_* represent the fitted rate of a gene (left-hand side) under the batch-free model described in equation 9; that is, let **μ**_i_* = E_Y_[**μ**_i_]. We finally obtain

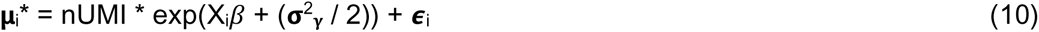

#### 1.4 Sampling corrected counts with inverse cumulative distribution function (iCDF) mapping

After obtaining an observed model and a batch-free model for a gene, we use these models to calculate a corrected count for each cell. Let x_i_ be the observed gene count and x_i_* be the corrected count for a cell *i*. Using the fitted GLMMs from the observed model and the batch-free model, we calculate fitted rates for each cell *i*: **μ**_i_ from the observed model (equation 3), and **μ**_i_* from the batch-free model (equation 10). We use **μ** and **μ**_i_* each to parameterize Poisson distributions for each cell *i*, which represent the probabilities of specific counts. We want to find a corrected count x* under the batch-free model (batch-free Poisson distribution) that has an equal probability of obtaining the observed count x under the observed model (observed Poisson distribution):

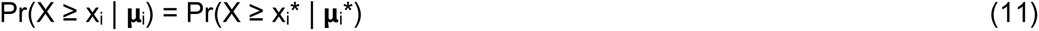

To calculate these probabilities, we calculate each distribution’s cumulative distribution function (CDF). With the CDFs of each distribution, fitted values for **μ**_i_ and **μ**_i_*, and the observed count x_i_ in a cell *i*, we want to find an x_i_* that satisfies equation 11. Let

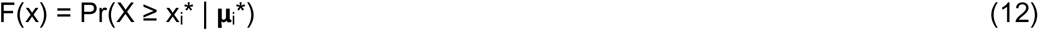

be the cumulative distribution function of the batch-free model. We use the inverse cumulative distribution function, or the quantile function, F^−1^(x*) to sample a corrected count x_i_* for each cell *i*. The quantile function returns a specific value that would have F(x) return a corresponding specific probability. We use this function to find a corrected count such that:

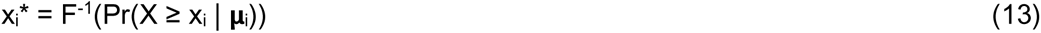

where x_i_* is the corrected count, x_i_ is the observed count, and **μ**_i_ is the fitted rate in the observed model from equation 3.

#### 1.5 Formalizing the Poisson framework as a mixture model

Crescendo corrects for cell-type specific batch effects. If all cell types were known *a priori*, we would simply apply the model above separately to each cell type. Importantly, we don’t assume perfect cell-type knowledge *a priori* and instead estimate latent variables that represent biological variation. These estimates are derived from the iterative soft clustering steps performed in Harmony. Because each cell is assigned probabilistically to each latent variable (i.e. soft clustering), we must model the effect of all cell types together. This is naturally formalized with a mixture model framework, similar to Harmony.

In this manuscript, we fit genes as mixture models that utilize the Harmony soft-cluster assignments (though we note that this framework is fully compatible with discrete cluster assignments). Fitting mixture models allows us to incorporate latent biological (cell-type) variation by fitting term coefficients for each cluster at the same time. Given K soft clusters, the GLMM in equation 3 is modified to have K beta (*β*) and K gamma (**γ**) terms. Conceptually, this is similar to fitting the wanted variation variables (X_i_) and the unwanted variation variables (Y_i_) for each cluster (though as previously stated, all of these terms are being fitted together due to soft cluster probabilities). The equation is modified to the following:

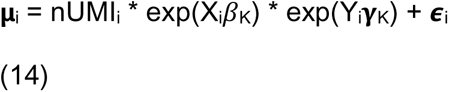

For the batch-free model in equation 10, we similarly incorporate the mixture model framework to give the following:

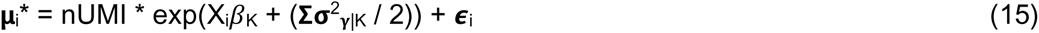

#### 1.6 Incorporating gene imputation with offsets

For imputation, the Poisson framework includes an offset representing exposure time, which is represented as the number of unique molecular identifiers (nUMIs) measured in a cell. This provides a natural framework to impute counts for a cell by fitting models in a context with higher-than-observed counts (e.g. sampling a cell’s corrected gene counts in a context of 10,000 UMIs).

The Poisson framework allows for the use of an offset term that represents exposure time, and is essentially fit as a fixed effect with a coefficient of 1. To fit the observed model in equation (3), we use the observed number of unique molecular identifiers (nUMI) of each cell for the offset. If users do not want to perform imputation in addition to correction, we also fit equation (10) with the observed nUMI.

Intuitively, we believe that it can be helpful to infer corrected counts for all cells in an equal UMI context, which allows for cell’s corrected counts to be inferred under equal exposure. That is, for calculating the final rates **μ**_i_ and **μ**_i_*, we use a constant nUMI instead of the observed nUMI. Note that for fitting the original GLMM, we still use the observed nUMI. Let cnUMI be a constant vector of length *i*. Then for imputation, equations 14 and 15 are modified as follows:

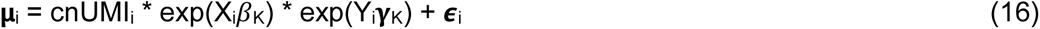

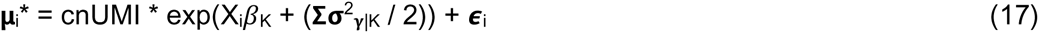

If users wish to impute at higher nUMI depths, then the constant value in cnUMI can be increased. For the results in this manuscript, we perform imputation under a constant nUMI context. The constant used is dataset-dependent, and is generally the median nUMI across all of the cells in the dataset.

#### 1.7 Batch and cell-type aware downsampling

Single-cell studies now often include more than 100,000 cells, while spatial transcriptomics datasets that include multiple slices may include millions of cells. While fitting count-based models for gene expression is more accurate than Gaussian models, they can take substantially longer to fit, especially as the dataset size increases. Furthermore, users may desire to fit more than one gene, which can mean fitting multiple models across millions of cells. To reduce the required computational resources and time for fitting, we allow users the option to downsample their data in a batch and cell-type aware manner. This downsampling is only for the purposes of fitting the GLMM for a gene, which constitutes the bulk of the computational runtime in Crescendo - all cells will be sampled corrected counts regardless of whether downsampling was utilized.

For downsampling in a batch and cell-type aware manner, we designate a minimum number of cells *m* so that we do not downsample too few cells. In this manuscript, we downsample the input dataset such that there are at least *m* cells within each cell-type within each batch in the downsampled dataset. If there are fewer than *m* cells within a cell-type within a batch, all cells of that type are kept. For more complicated data structures such as nested batch structures, we suggest downsampling such that the lowest-level groups have at least *m* cells; these would require a custom downsampling function depending on the data structure. By default, we utilize Harmony soft-cluster assignments, which means that a cell may have membership in multiple clusters. For the purposes of downsampling, we assign each cell a discrete cell-type label by creating a probability distribution from its soft-cluster membership probabilities, and then we sample a cluster label from this distribution.

We allow users to specify a proportion, which proportionally downsamples the number of cells within a cell-type within a batch (e.g. a proportion of 0.25 will try to sample 25% of cells in a cell-type in a batch, unless there are fewer than *m* such cells). In general, we tended to fit on around 20,000 cells total for each dataset. By default, we set *m* = 100, but it is likely that fewer cells are required to obtain relatively similar coefficients to the full dataset.

### 2. Calculating batch-variance ratios and cell-type variance ratios for performance metrics

In scRNA-seq, the performance of an integration algorithm is often evaluated by how they change the structure of the data in a low-dimensional latent space. Typically, integration algorithms increase the diversity of batches in a local area of the latent space, which is quantified with a metric. Because these latent spaces are summarizations of many genes, we cannot directly apply previously created metrics to quantifying integration performance in a single gene. Thus, we now describe two metrics which can be used to evaluate the performance of integration in genes.

Effective batch effect correction of gene expression must meet two objectives: (1) remove differences (variation) between cells of the same cell type that are driven by technical factors such as batch and (2) preserve the biologically meaningful differences in gene expression among cell types. With these objectives in mind, we developed two metrics that each address one of these objectives. The first metric, which we call the batch-variance ratio (BVR), quantifies how much batch effect was removed from a gene count distribution after correction, while the second metric, cell-type-variance ratio (CVR) quantifies the preservation of cell-type variation after correction.

To calculate the BVR and CVR metrics, we fit Poisson generalized linear models (GLMs) that estimate the batch variance and the cell-type variance present in a given count distribution. We calculate these variances by fitting batch and cell-type as independent random effects, as well as an independent interaction term between batch and cell-type to estimate cell-type-specific batch variance. In practice, we utilize user-defined discrete clusters (e.g. T cell, B cell). To fit Poisson GLMs, we used the R package “presto”, which utilizes the “glmer” function from the R package “lme4”. For the the observed counts X, we fit the following formula:

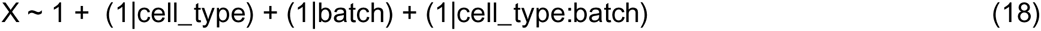

For the corrected counts X*, we similarly fit:

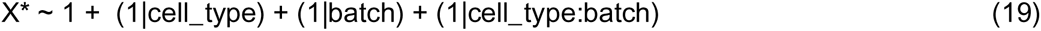

For fitting the Poisson GLMs in equation 18 and 19, we use the observed nUMI for cells as the offset.

To calculate the BVR, we obtain the variance estimates for the batch, and cell-type-specific batch terms. For simplicity, we calculate the overall batch variance estimate as the sum of the cell-type-specific batch and batch estimates. We obtain an overall batch-variance estimate from the corrected counts based on equation 19 in the same way. Let B_pre_ be the pre-correction batch-variance estimate obtained from equation 18 and let B_post_ be the post-correction batch-variance estimate obtained from equation 19. We calculate BVR as:

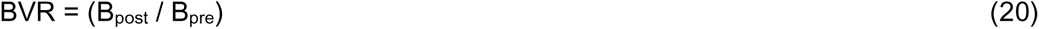

In a similar manner, we obtain cell-type variance estimates from both equation 18 and 19. Let C_pre_ be the pre-correction cell-type-variance estimate from equation 18 and let C_post_ be the post-correction cell-type-variance estimate from equation 19. We calculate CVR as:

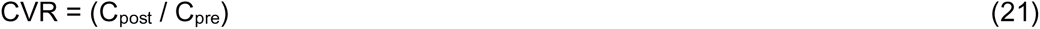

The BVR metric quantifies how much batch-related variance was removed after correction, while the CVR metric quantifies how much cell-type-related variance was preserved after correction. Based on the two objectives we outlined at the beginning of this section, ideal correction will decrease batch variance resulting in a BVR < 1, while preserving or increasing cell-type variance resulting in a CVR >= 1. In practice, integration usually features a trade-off - the more aggressively batch effects are removed, the more cell-type variance tends to be removed (although sometimes cell-type variance is also increased if a gene becomes more specific to a cell-type after correction). Empirically, a CVR >= 0.5 was a reasonable trade-off if the BVR was lowered.

We also note that the batch-related variance value before correction may be a useful value for users, as it can help determine which genes have higher levels of batch effects and might need correction.

### 3. Gene count simulations

We simulated gene count distributions by sampling from Poisson distributions parameterized by different rates based on the cell-type or batch a cell is from. To simulate a single gene, we designate the number of batches, as well as the number of cells we will simulate for each cell type per batch. Each cell belongs to one cell type and one batch. We then arbitrarily set a base rate for each cell type (e.g. a rate of 1 for cell-type 1 and a rate of 3 for cell-type 2). To simulate batches, we sampled a batch-specific rate for each batch from a standard normal distribution, and then centered all batch-specific rates around 0. For simplicity and visualization, we simulated from two cell types in two batches, though this framework is compatible with an arbitrary number of cell types and batches. After sampling batch-specific rates, we add them to the base rate for each cell type. for example, batch 1 will add a batch-specific rate of 0.405 to cell-type 1’s rate of 1 to result in a unique rate of 1.405, while batch 2 will add a batch-specific rate of −0.405 to cell-type 1’s rate of 1 to result in a unique rate of 0.595. Thus, each cell type within each batch has its own unique rate that represents a batch effect.

We then assign each cell membership a probability membership for each cell type, which represents soft-cluster membership. For two cell types, we sampled from a beta distribution parameterized with ɑ = 0.5, β = 0.5 to get the probability *p* of a cell belonging to one cell-type; to calculate the probability *q* or a cell belonging to the other cell-type, we simply take 1-*p*. We also set each cell to have the same constant number of unique molecular identifiers (nUMI), though this framework is compatible with variable nUMIs (recommend sampling from a lognormal distribution).

We then created a design matrix that contains the cell-type probabilities and the batch identities of each cell, and then matrix-multiplied the design matrix with a matrix containing the batch-specific rates for each cell type. After, we multiplied the resulting product with a matrix containing the cell-type probabilities to recover a rate for each cell based on its batch and cell-type identity. To represent read depth, we add a log-transformed nUMI constant to each cell’s rate (in our simulations, we set the constant for each cell to be equal at 10,000). Finally, we use the resulting rates to parameterize a Poisson distribution for each cell, which we then sample a count from.

For **Supplementary Figure 1F**, we simulated 10,000 genes, with the base cell-type rates set at 1 for cell-type 1 and 3 for cell-type 2.

### 4. Plotting gene expression visualizations

To plot gene expression across batches, we utilize the “facet_wrap” function from the R package “ggplot2”. This function allows us to visualize the same gene’s expression across all batches together on the same scale. For visualization purposes, we plot cells that express a gene on top of other cells that don’t express the gene. This represents a best-case scenario in which we should be able to see every instance of gene expression, and is extremely forgiving if the gene is poorly expressed. In practice, most visualization of data is performed with cells being randomly mixed such that gene-expressing cells are not always on top. In such scenarios, we observed that Crescendo dramatically improves visualization even more than the best-case scenario, which is already a significant improvement.

### 5. Benchmarking and comparison to other algorithms

To perform benchmarking in **Figure 2 and Figure 3**, we compared Crescendo with the following algorithms: ComBat-Seq, Seurat anchor integration, limma, and mutual nearest-neighbors (MNN) correction.

To perform integration with ComBat-Seq, we used the “ComBat_seq” function from the R Bioconductor package “sva” with default parameters. ComBat-Seq is designed to fit and output counts, so we calculated BVR and CVR metrics based on fitting Poisson models of the raw gene expression counts and the ComBat-Seq corrected gene expression counts (**Section 2**).

For integration with Seurat, we used Seurat version 4.3.0 in R. For each batch, we created a Seurat object and normalized them with the “NormalizeData” function. We then integrated the datasets with the “FindIntegrationAnchors” and “IntegrateData” functions with default parameters and dims = 1:20. To access corrected counts from the integration, we accessed the object’s “@assays$integrated@data” slot. Seurat’s integration works on normalized gene expression and returns gene expression in a similar normalized space, so we calculated BVR and CVR metrics based on fitting Gaussian models of the normalized gene expression counts and the Seurat-corrected counts (**Section 2**).

For limma, we used the “removeBatchEffect” function from the R package “limma” with default parameters. Limma’s integration works on normalized gene expression and returns gene expression in a similar normalized space, so we calculated BVR and CVR metrics based on fitting Gaussian models of the normalized gene expression counts and the limma-corrected counts (**Section 2**).

For MNN, we used the “fastMNN” function from the R package “batchelor” with default parameters. MNN’s integration works on cosine-normalized gene expression and returns gene expression in a similar normalized space, so we calculated BVR and CVR metrics based on fitting Gaussian models of the cosine-normalized gene expression counts and the MNN-corrected counts (**Section 2**).

### 5. Vizgen Mouse Brain Receptor analysis details

We downloaded the Vizgen Mouse Brain Receptor metadata and count matrices from the Vizgen Data Release Program. This dataset contains a panel of 483 genes. For **Figure 2**, we subsetted the data to only include cells from slice 3 (S3R1, S3R2, S3R3), which represent serial sections from the same mouse brain (186,910 total cells). Following Vizgen recommendations, we filtered out cells with fewer than 50 total expressed transcripts or fewer than 50 uniquely expressed genes, resulting in 179,385 remaining cells: 53,269 from S3R1, 64,476 from S3R2, and 61,640 from S3R3. For the following steps, we used all 483 genes. We library-normalized cells with standard llog-normalization with the median read counts as the scale factor, and scaled genes with z-score scaling. We then utilized PCA to reduce the dimensionality of the data to the top 20 PCs, and performed integration with the Harmony algorithm. To cluster cells, we utilized Leiden clustering with resolution = 0.2. Finally, we used Crescendo to correct all genes using S3R1, S3R2, and S3R3 as batches. We used the observed nUMI as the initial offset, and then used the median nUMI as the final offset for imputation. For visualization in physical space, we rotated each slice’s coordinates such that they are in the same orientation.

### 6. Scalability analysis

For the scalability analyses, we utilized the public Vizgen FFPE Immuno-oncology dataset. We downloaded the metadata and count matrices from the Vizgen Data Release Program. This dataset contains a panel of 500 genes measured on 16 human cancer samples across 9 different tissue types (∼8.7M total cells). Following Vizgen recommendations, we filtered out cells with fewer than 50 total expressed transcripts or fewer than 50 uniquely expressed genes, resulting in 7,020,548 remaining cells. We library-normalized cells with standard log-normalization with the median read counts as the scale factor, and scaled genes with z-score scaling. We then utilized PCA to reduce the dimensionality of the data to the top 20 PCs and performed integration the Harmony algorithm. To correct with Crescendo, we used sample identity (Lung Sample 1, Lung Sample 2, Liver Sample 1, etc.) as batches. We used the observed nUMI as the initial offset, and then used the median nUMI as the final offset for imputation.

To accommodate the large memory required to load this dataset, integration and scalability analyses on this dataset were run on a server containing 24 cores and 128GB of RAM for all algorithms.

### 7. Integrated colorectal cancer (CRC) scRNA-seq and spatial transcriptomics analysis

For the CRC scRNA-seq dataset, we downloaded the metadata and count matrices from GEO: https://www.ncbi.nlm.nih.gov/geo/query/acc.cgi?acc=GSE178341. We first filtered for only cells from SPECIMEN_TYPE = T (cells taken from tumor samples) and SINGLECELL_TYPE = SC3Pv3 (only cells assayed with 10X v3), which resulted in 90,312 remaining cells. We further QCed to keep only cells that featured a total nUMI from 30-2000 counts and expression in at least 10 unique genes, resulting in 86,627 cells. Spatial samples were taken from one of the donors in this dataset – we kept all donors in this datatet as the spatial donor only had ∼1,600 scRNA-seq cells.

The spatial transcriptomics tissues were produced in collaboration with Vizgen. These tissues derive from the same patient sample, which is also represented in the scRNA-seq data. Segmentation was performed with Baysor.

After combining scRNA-seq and spatial data, we library-normalized cells with standard log-normalization with the median read counts as the scale factor, and scaled genes with z-score scaling. We then utilized PCA to reduce the dimensionality of the data to the top 20 PCs and performed integration with the Harmony algorithm. To cluster cells, we utilized Leiden clustering with resolution = 0.1. To perform correction with Crescendo, we represented scRNA-seq as its own batch and the two spatial transcriptomics slices as their own individual batch (scRNA-seq, PFA_A6, and PFA_A11 were the batches). We used the observed nUMI as the initial offset, and then used the median nUMI as the final offset for imputation.

### 8. Spatial cross-correlation index (SCI) calculations

To calculate an SCI in a cell-type-aware manner, we first subsetted a spatial transcriptomic dataset’s count matrix to two (user-specified) cell types. We then calculated the 30 nearest-neighbors for each cell (excluding itself) with the “nn2” function from the R package “RANN” and retrieved a sparse distance matrix from the nn2 output with the “getDistMat” function provided by Crescendo. We next removed a cell’s neighbors if they are the same cell-type (by setting its value to 0 in the distance matrix). We then removed neighbors with a distance > 30μm from the cell. Finally, we binarized the matrix by setting all non-zero values to 1. This binarized matrix (K) contains information on whether another cell is a nearest-neighbor, a different cell type, and within a distance of 30. Thus, for a cell-type 1, we have its nearest-neighbors from cell-type 2 and vice-versa.

We then take the subsetted raw gene count matrix and log-normalize the counts with the median nUMI as the scale factor to produce a normalized gene counts matrix X. Then, we matrix-multiplied the raw gene count matrix with the binarized matrix (K) to produce a normalized gene nearest-neighbors gene counts matrix (XK). Thus, X contains the gene expression of cells while XK contains the gene expression of that cell’s nearest neighbors from the other cell type. Finally, we use the “cor” function from base R with X and XK as input and with default parameters to obtain the correlations for each gene-gene pair. SCI calculations were performed in each slice independently.

Two genes that share similar spatial expression patterns will exhibit a higher SCI, while two genes whose spatial patterns are not correlated with exhibit a low SCI.

## Data availability

All datasets except for the colorectal cancer (CRC) spatial transcriptomics tissues are publicly available through online sources. CRC spatial tissue data will be made publicly available in a future publication.

## Code availability

Crescendo will be made available as an R package on https://github.com/immunogenomics/crescendo, along with a vignette to correct gene expression.

## Acknowledgments and Attributions

## Acknowledgments

We thank members of the Hacohen lab and the Vizgen research team for their contributions to generating the colorectal cancer (CRC) scRNA-seq and spatial transcriptomics datasets.

## Author Contributions

N.M, I.K, and S.R. conceived the project. N. M. developed the method and performed the analyses under the guidance of S.R. and I.K. J.C. and P.K. generated the CRC scRNA-seq dataset. M.G.P. provided computational analysis of single-cell datasets. J.C. M.S., K.P., J.H. and C.P. designed the CRC spatial transcriptomics panel, constructed the cohort, and generated spatial transcriptomics datasets. All authors participated in the interpretation and writing of the manuscript.

