## Supplementary Figures for "Integrating spatial transcriptomics count data with Crescendo improves visualization and detection of spatial gene patterns"

### Supplementary Figure 1

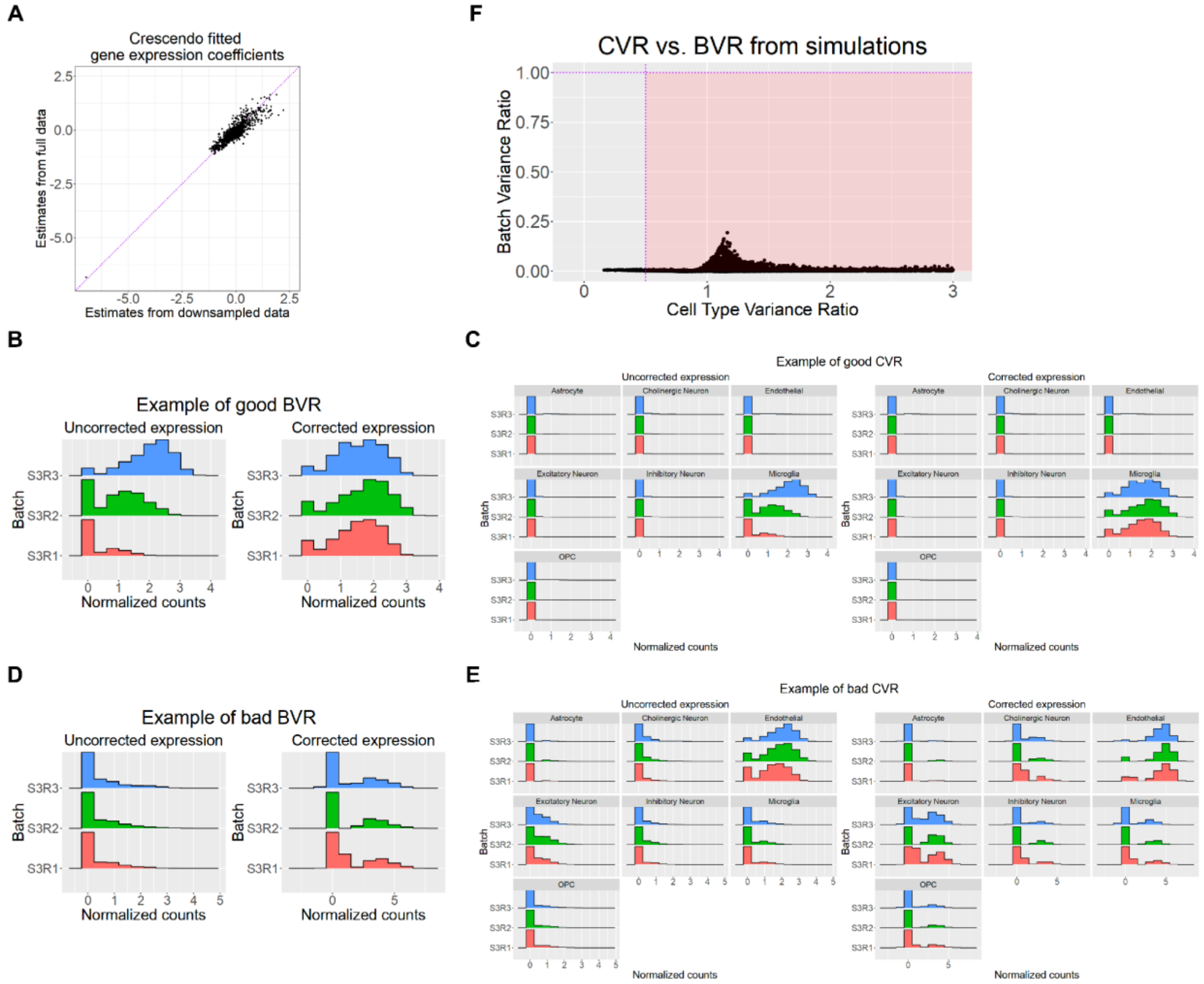

**Supplementary Figure 2**

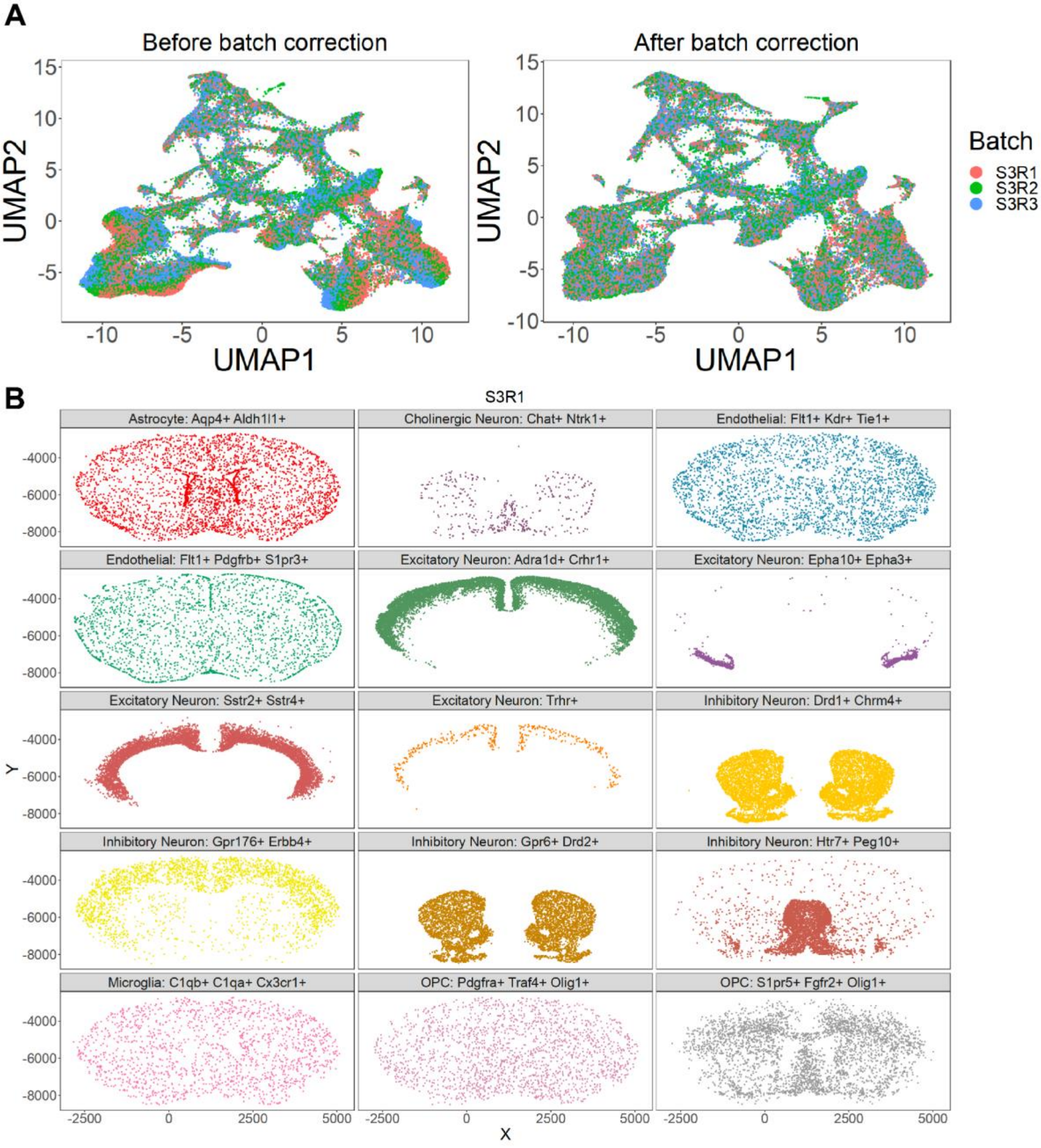

**Supplementary Figure 3**

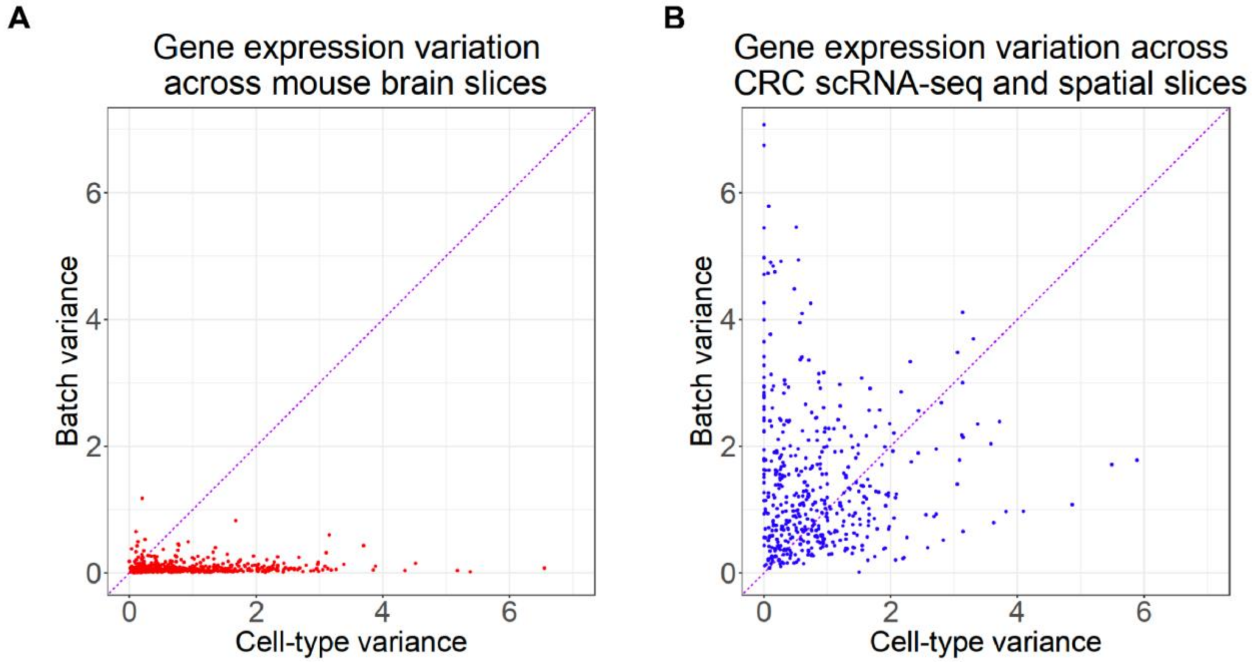

**Supplementary Figure 4**

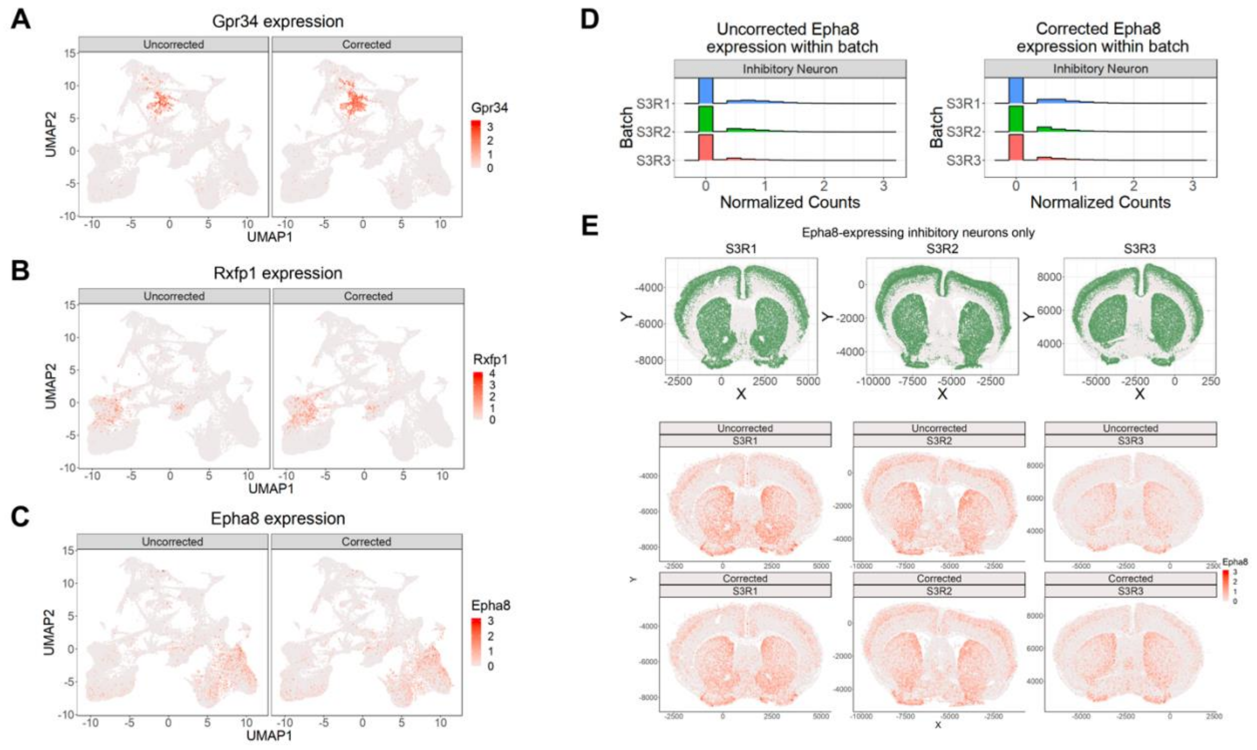

**Supplementary Figure 5**

**A**

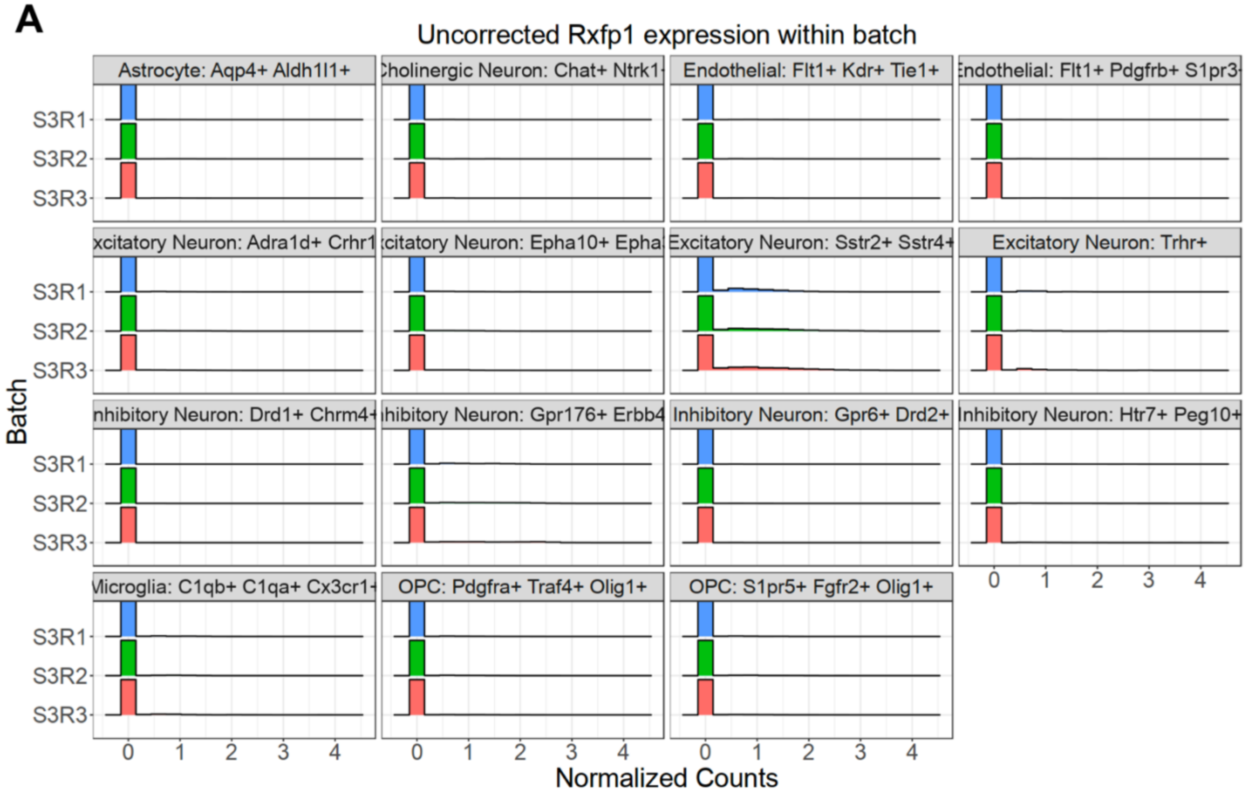


**B**

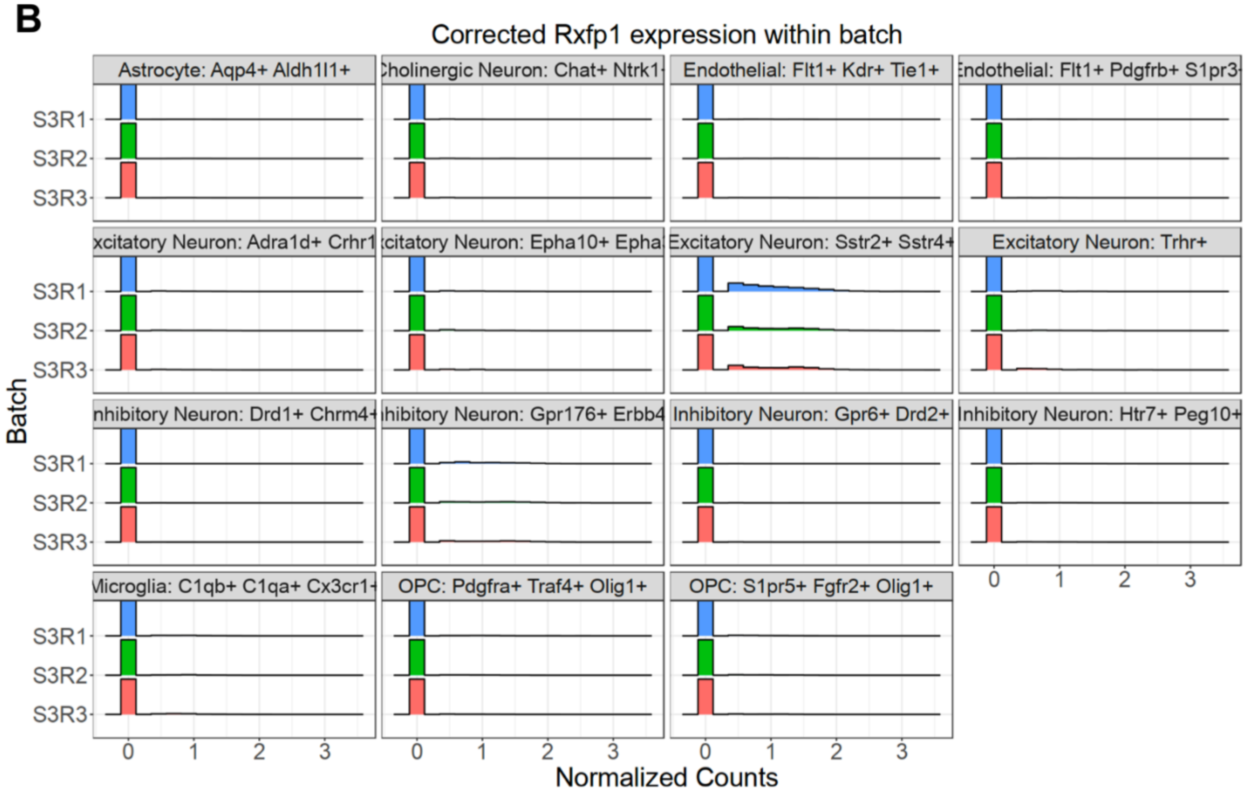

**Supplementary Figure 6**

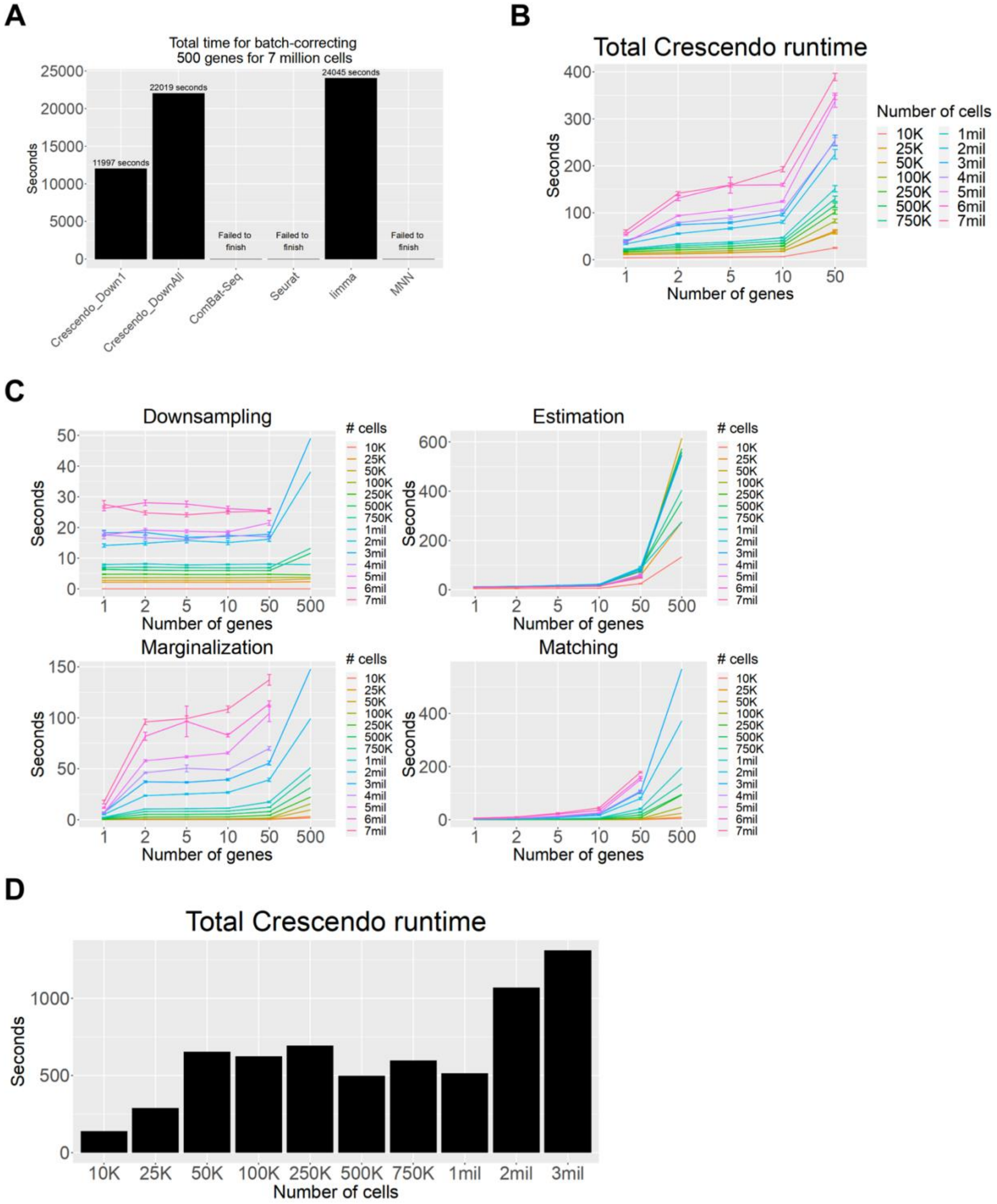

**Supplementary Figure 7**

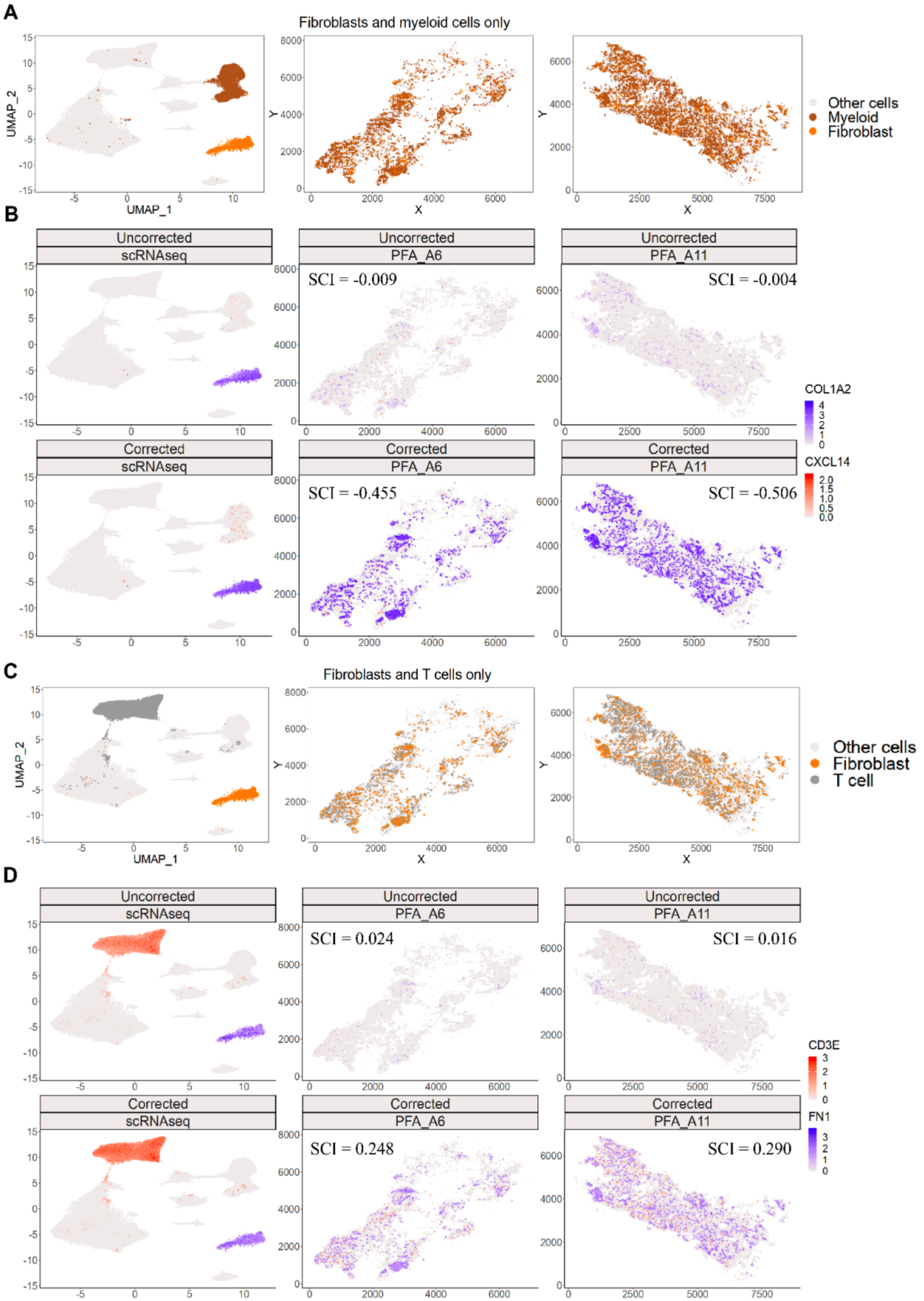
